# Avoiding Math on a Rapid Timescale: Emotional Responsivity and Anxious Attention in Math Anxiety

**DOI:** 10.1101/131433

**Authors:** Rachel G. Pizzie, David J.M. Kraemer

## Abstract

Math anxiety (MA) is characterized by negative feelings towards mathematics, resulting in avoidance of math classes and of careers that rely on mathematical skills. Focused on a long timescale, this research may miss important cognitive and affective processes that operate moment-to-moment, changing rapid reactions even when a student simply sees a math problem. Here, using fMRI with an attentional deployment paradigm, we show that MA influences rapid spontaneous emotional and attentional responses to mathematical stimuli upon brief presentation. Critically, participants viewed but did not attempt to solve the problems. Indicating increased threat reactivity to even brief presentations of math problems, increased MA was associated with increased amygdala response during math viewing trials. Functionally and anatomically defined amygdala ROIs yielded similar results, indicating robustness of the finding. Similar to the pattern of vigilance and avoidance observed in specific phobia, behavioral results of the attentional paradigm demonstrated that MA is associated with attentional disengagement for mathematical symbols. This attentional avoidance is specific to math stimuli; when viewing negatively-valenced images, MA is correlated with attentional engagement, similar to other forms of anxiety. These results indicate that even brief exposure to mathematics triggers a neural response related to threat avoidance in highly MA individuals.

## Avoiding Math on a Rapid Timescale

### Emotional Responsivity and Attentional Aversion in Math Anxiety

A major challenge to students’ interest and success in STEM courses is math anxiety, or negative and anxious emotion associated with anticipation and performance of mathematics (Ashcraft, 2002; Hembree, 1990). While much research has focused on the long-term outcomes of math anxiety, less work has examined the low-level rapid or automatic responses that occur each time a highly math anxious (HMA) individual encounters math stimuli. This is an important consideration for understanding the neural, cognitive, and affective mechanisms underlying math anxiety, and ultimately, for treating these negative outcomes. For instance, suppose math anxiety is associated with an initial, rapid aversion to seeing a problem on a math test, even before beginning to solve the problem at hand. This type of reaction represents an obstacle in terms of the resources required for a HMA student to even begin the process of computing the solution to the problem. Prior to engaging the neural and cognitive resources that are typically brought to the task, a math anxious student must first respond to – and effectively inhibit – the threat signal that his or her emotional response system is generating. In the present fMRI experiment, we begin to examine the dynamics of how this process unfolds over a rapid timescale – on both a behavioral and a neural level – which will inform our understanding of how math anxiety influences cognition.

Neuroimaging presents a valuable tool to better understand how negative affect can alter initial neural reactivity to math, and how this emotion ultimately alters mathematical computation in the brains of HMA individuals. The amygdala responds to a myriad of affective cues, implying that this area of the brain is not just selective for fearful facial expressions (Whalen, 1998; 2004b), but instead provides a mechanism for attentional mechanisms for uncertainty and resulting in cortical vigilance (Davis & Whalen, 2001). We examine patterns of neural reactivity in the amygdala because math anxiety may be a result of learned negative responses, much like fear conditioning, such that mathematics may be thought of as a negatively conditioned stimulus. In generating affective responses in learning and fear conditioning (Davis & Whalen, 2001; Holland & Gallagher, 1999), the amygdala associates information linking a conditioned stimulus to positive or negative outcomes, signaling this value to the nucleus accumbens (NAcc) in the ventral striatum. In turn, dopaminergic projections from the NAcc back to the amygdala amplify this signal in a feedback loop. Additionally, the amygdala is responsible for lowering neuronal thresholds in sensory systems, through activation of basal forebrain cholinergic neurons, serotonergic systems and catecholamnergic systems, thereby increasing vigilance in order to facilitate a response during uncertain or potentially threatening situations (Davis & Whalen, 2001).

This link between increased amygdala reactivity and a lowered threshold for responding to potentially threatening cues is a hallmark of many types of anxiety and phobia. Anxious individuals show increased amygdala reactivity to negative information and increased attentional bias for negative cues, as this threshold for attending and reacting to negative information is lowered (Bishop, 2007; 2008). Similarly, individuals with specific phobia show increased amygdala reactivity when viewing phobic stimuli (Schienle, Schäfer, Walter, Stark, & Vaitl, 2005), illustrating increased vigilance for a stimulus associated with a specific learned fearful reaction. However, unlike the increased attentional engagement observed for anxious individuals, phobia is associated with behavioral avoidance of phobic stimuli (Pflugshaupt et al., 2005). In this way, HMA individuals may have developed these anxious and negative responses to mathematics much in the way that someone learns to respond to a conditioned stimulus. This threat system may be triggered just by viewing a math problem, before other regions of the brain associated with mathematical processing (e.g., the intraparietal sulcus, IPS; Dehaene, Spelke, Pinel, Stanescu, & Tsivkin, 1999) are engaged to complete these computations. In other words, the negative affective response in math anxiety may begin very rapidly, even with mere exposure to a math problem, and thus alter subsequent attention, and further downstream, could alter the neural basis of mathematical cognition when individuals are asked to complete mathematical computations. In the present fMRI experiment, we test whether math anxiety is associated with initial hypervigilance and negative reactivity in the amygdala, illustrating that math anxiety alters even early responsivity to mathematical information, even before one begins the process of computation.

In line with the description of math anxiety as a tradeoff in neural resources between negative emotion processing and cognitive operations pertaining to math, previous research indicates HMA individuals show increased amygdala reactivity while performing math computations, and decreased activity in regions associated with math computation (Young, Wu, & Menon, 2012). However, it is possible that – even without requiring completion of math computations – increased affective processing for HMA participants results in specific arousal-related amygdala reactivity. This possibility has been left unexplored by previous work, which has focused either on periods of math computation (Young et al., 2012), or anxious anticipation of math computation (Lyons & Beilock, 2012a; 2012b). In contrast, here we examine responsivity to brief presentation of math stimuli absent any computation or instruction to evaluate numerical values. Therefore, if the results of this experiment demonstrate that amygdala responsivity while attending to math stimuli varies as a function of math anxiety, we can conclude that this response is not due to anticipating or engaging in an unpleasant mathematical task, but simply to seeing a math problem. In this way, due to repeated negative experiences, math problems may become a negative conditioned stimulus, and even brief exposure to these stimuli may evoke a rapid negative reaction. In other words, these results would suggest that negative amygdala reactivity in math anxiety cannot be wholly attributed to directly experiencing or anxiously awaiting a subjectively negative task. Instead, this increased emotional arousal and vigilance in response to a negatively-valenced stimulus occurs on a very rapid timescale, shaping initial attention and perhaps altering deployment of cognitive resources that could affect later working memory and mathematical computation.

In the present fMRI study, we explored how math anxiety influences neural and behavioral indices of anxious emotion and rapid orienting responses to math stimuli, illustrating heightened negative reactivity even without solving math problems. We utilized a dot probe task in an fMRI experiment examining how math anxiety influences attentional responses to briefly-presented mathematical expressions, as a means of measuring rapid emotional and attentional responsivity. By establishing the attentional changes and neural patterns that are associated with brief exposure to mathematical stimuli, we aim to develop a greater understanding of how negative emotion manifests in math anxiety and influences mathematical cognition and understanding.

## Method

### Power Analyses

Using the effect size from a previous dot probe study with anxious participants, (Cohen’s *d = .79, r*^2^ = .135*, f* = .391; Bradley, Mogg, Falla, & Hamilton, 1998), for 80% power at α *=* .05, we would require *N* = 36 participants, so 40 participants were recruited.

### Participants

Forty undergraduate students completed a dot probe attentional paradigm in the fMRI scanner. Participants were recruited from a subject pool on the basis of their extreme scores on the Math Anxiety Rating Scale, (MARS; Suinn & Winston, 2003). The resulting sample included 20 high math anxious individuals (HMA, *M*_*MARS*_= 2.98, *SD*_*MARS*_ = .21, *M*_*age*_ = 19.30 years, 70% female) and 20 low math anxious individuals (LMA, M_MARS_ = 1.60, *SD*_*MARS*_ = .21, *M*_*age*_=19.65 years, 55% female).

### Task

This fMRI task was designed to assess attentional deployment to traditionally negative stimuli (IAPS pictures; Lang & Bradley, 2007), as well as mathematical expressions. In the dot probe paradigm, attention to a stimulus of interest is compared to a neutral stimulus by measuring the speed and accuracy of responses to a subsequent stimulus in the same or different spatial location (MacLeod & Mathews, 1988). If participants are comparatively faster to detect the appearance of a dot probe in the same location as the stimulus of interest, this indicates an engagement bias towards the initial stimulus. Conversely, individuals show a disengagement bias to the degree that they show improved performance when the dot probe appeared in the opposite location as the stimulus of interest, indicating that they directed attention away from this stimulus. If math anxiety influences attention in a similar manner to trait anxiety, we would expect to observe an engagement bias for mathematical stimuli for HMA individuals (MacLeod & Mathews, 1988). However, previous work on math anxiety also suggests that HMA individuals tend to avoid mathematics (Ashcraft, 2002). If, like specific phobia (Pflugshaupt et al., 2005), these broader patterns of avoidance behavior are associated with rapid, compulsory avoidance of specific stimuli upon brief presentation, response times will reveal a disengagement bias instead.

Each trial was composed of 1000 ms of image/symbol stimulus presentation, 1000 ms of dot probe presentation (250 ms) and detection (750 ms), and 500 ms of fixation, such that each trial lasted for a total of 2500 ms (Figure 1). Inter-trial-intervals of jittered fixation between 0 ms and 12500 ms occurred after each trial. The stimuli presented were either images or symbols, thus allowing us to compare allocation of attention to negative images (“negative” condition) compared to control neutral images (“neutral”); and mathematical symbols (“math”) could be compared to control linguistic symbols (“symbol”). Participants were tested over 3 blocks of trials, totaling 60 of each trial type, for a total of 240 trials overall.

**Figure 1.**
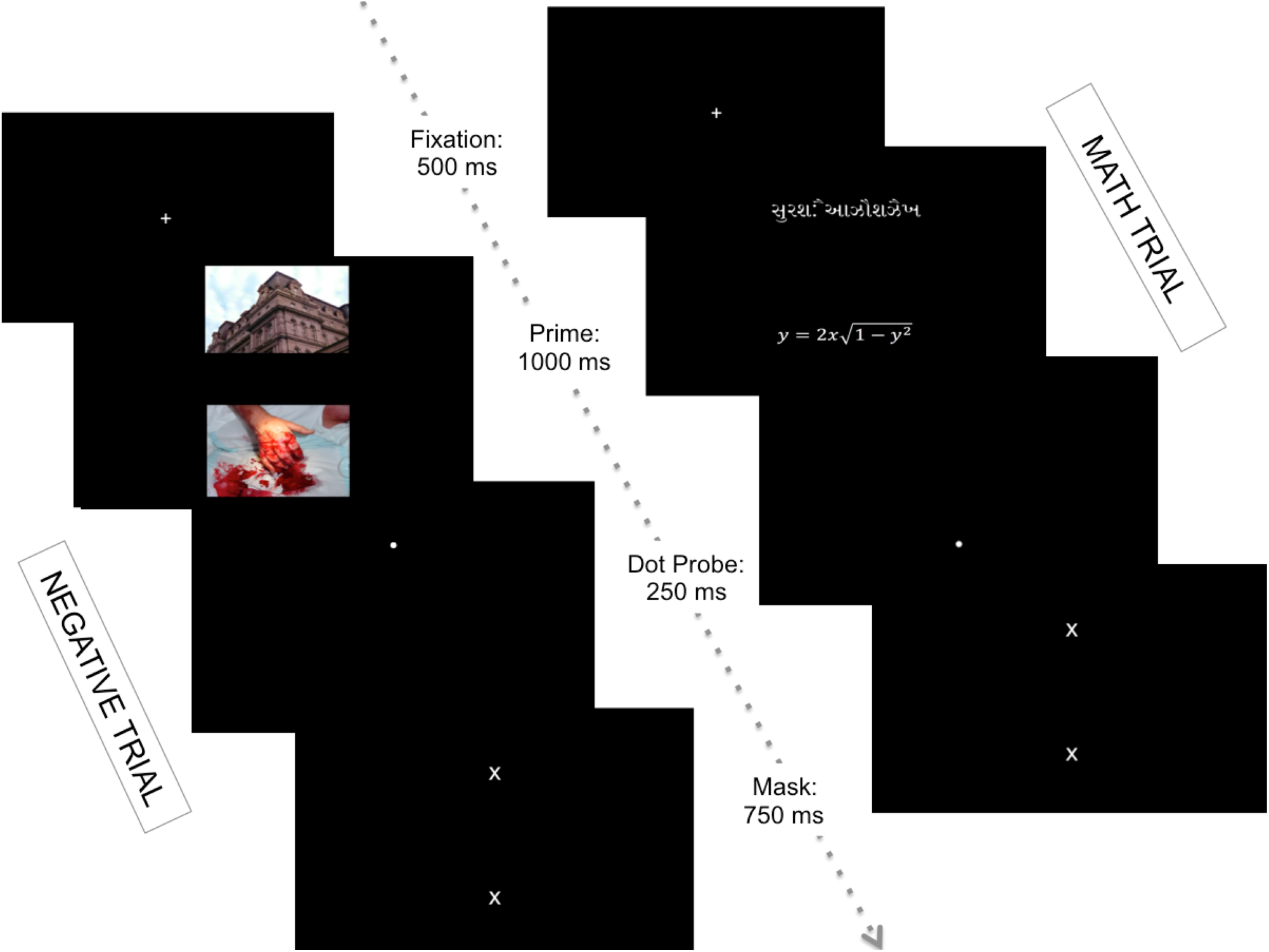
Trial Cadence for Dot Probe Task. Note: Examples of negative (left) and math (right) trials are pictured. Participants are told to indicate the location of the dot probe with a button press. Response times and accuracy are recorded from the onset of the dot probe for the duration of the mask (a total of 1000 ms). The math trial represents a “congruent” trial as both the math stimulus and dot probe appear in the same spatial location. The negative trial depicts an “incongruent” trial as the stimulus of interest (negative picture) and dot probe appear in opposite locations. Equal numbers of trials were presented in each condition (60 each for math, symbol, negative, neutral) for a total of 240 trials.

After scanning was complete, participants completed assessments of trait anxiety (Spielberger State-Trait Anxiety Inventory—trait, STAI; Spielberger, 2010), test anxiety (Spielberger Test Anxiety Inventory, TAI; Spielberger, 2009), writing anxiety (WA; Daly & Wilson, 1983), assessments of positive and negative affect (PANAS; Watson, Clark, & Tellegen, 1988), and provided demographic information. One participant was missing a STAI score and was excluded from analyses examining trait anxiety. Outliers (scores ± 3 standard deviations from the mean) for each questionnaire were excluded from the relevant analysis. After completing these questionnaires, participants were debriefed and thanked for their participation.

### Image Acquisition

Participants performed this task in the fMRI scanner using a rapid event-related design. Functional scans used a 80 x 80 reconstruction matrix in a 240 mm^2^ FOV for whole-brain coverage (Flip angle= 90 degrees, TE = 35 ms, TR = 2500 ms, 3 mm^3^ voxels). For each of 3 functional runs, 170 TRs of trials and fixation were collected, for a total of 510 functional volumes.

### ROI Selection

We identified *a priori* regions of interest (ROIs) specifically associated with processing of negative affect, and previous research has indicated the amygdala as a region of the brain associated with orienting to negative information (Whalen, 1998), and increased attention to negative information is exaggerated in increased anxiety (Bishop, 2008). We identified functional ROIs in left and right amygdala based on the contrast of Negative > Neutral images in a whole-brain GLM. A sphere of voxels (4 mm radius) centered at the peak of activation in the left amygdala (*t*_*peak*_ = 5.37, MNI coordinates: *x* = -22, *y* = 4, *z* = -20), and right amygdala (*t*_*peak*_ = 4.42, MNI coordinates: *x* = 28, *y* = -4, *z* = 22) were each used as ROIs (for all clusters from the Negative > Neutral contrast, see Supplemental Table 1). We calculated signal change within each amygdala ROI for the Math > Symbol contrast for each individual participant. To examine the consistency of this relationship, we also examined functional activity in the amygdala extracted from an anatomically defined ROI, and a whole-brain GLM analysis to determine whether activity in other regions of the brain demonstrates this same correlation with math anxiety. For analyses using anatomically-defined ROIs, FreeSurfer cortical and subcortical parcellations were used to create masks of the amygdala for each subject (Dale, Fischl, & Sereno, 1999). Using left and right anatomical amygdala masks, we calculated signal change within each anatomical ROI for the Math > Symbol contrast for each participant. (For additional information on behavioral and neuroimaging analyses, see Supplementary Materials).

## Results

### Behavioral Results

Accuracy scores for the dot probe task showed ceiling effects, with average accuracy across all trials was close to 100% (*M* = 94.1% correct, *SD* = 11.9). However, one participant was dropped from further behavioral analyses for low accuracy across conditions and below-chance responding (*M* = 27% correct trials, chance level = 50%, > 3 SD away from mean). The resulting sample included 39 subjects (*M* = 95.9% correct, *SD* = 4.92). Only correct trials were used for further analysis of response time (RT). Outliers (> 3 SD away from mean) on survey scores or response times were excluded from the relevant analyses.

To investigate this relationship with math anxiety scores, we conducted a mixed ANOVA to determine whether these engagement/disengagement bias scores differed based on condition (within-subject factor, 2: math, negative) and individual differences in level of math anxiety (MARS score: between-subject factor). A significant interaction was found for bias scores based on individual level of math anxiety, such that HMA individuals show an engagement bias for negative trials, and a disengagement bias for math trials, *F*(1,36) = 5.86, *p* = .02, *η*^2^ = .08 (Figure 2). However, the range of these scores is restricted to high and low scores, and does not represent the full range of math anxiety. When participants were grouped by HMA vs. LMA, HMA individuals show an engagement bias for negative trials, and a slight disengagement bias for math trials, *F*(1,37) = 3.61, *p* = .065, *η*^2^ = .05, though this relationship is not statistically significant.

**Figure 2.**
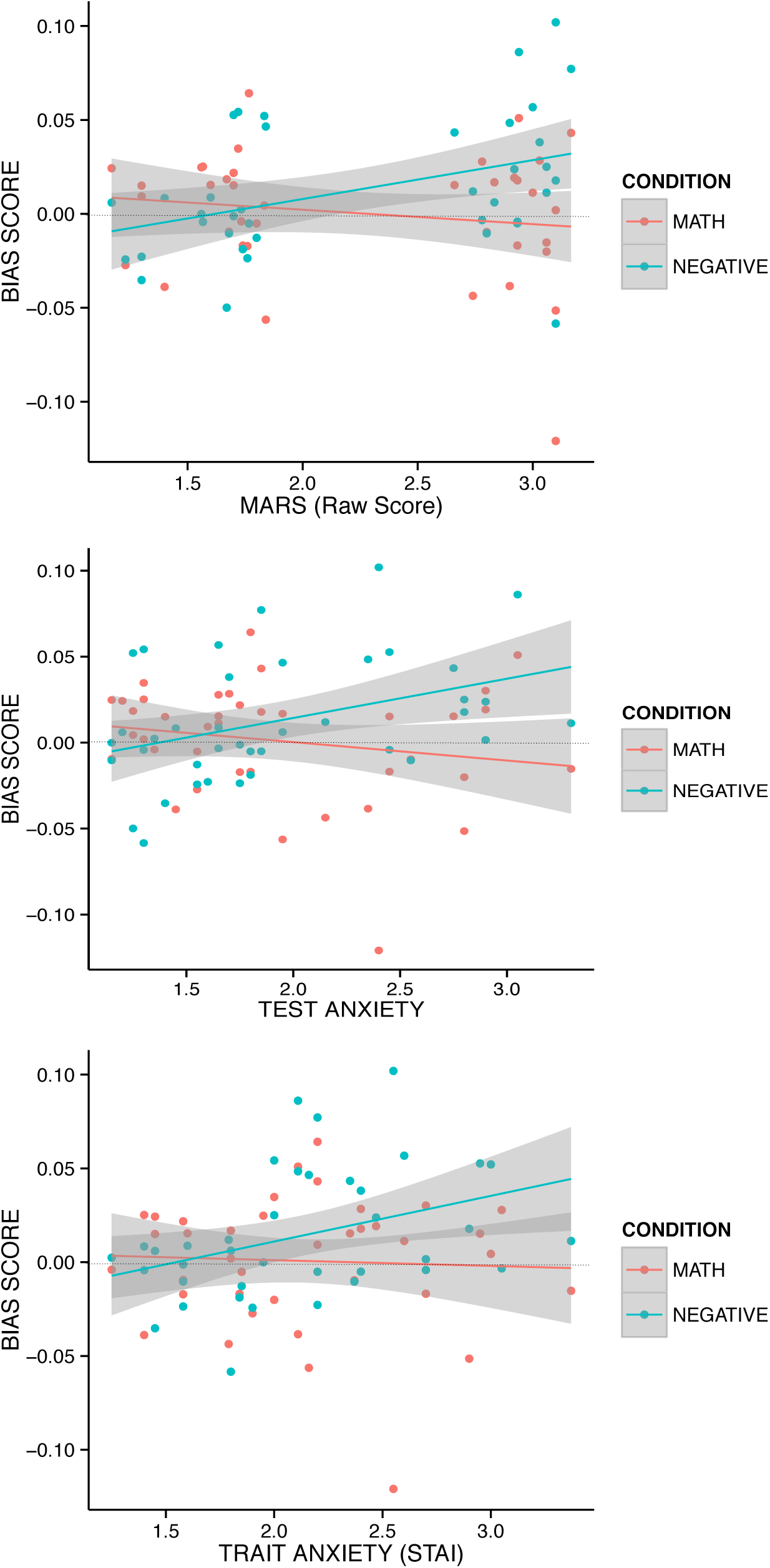
Engagement and Disengagement bias scores. Note: Bias score is calculated by subtracting Incongruent – Congruent response times from the baseline-controlled scores negative and math scores (negative and neutral response times are subtracted from the respective experimental condition). Positive scores indicate an engagement bias. Negative scores indicate a disengagement bias. Outliers (> 3 SD away from mean) on survey scores or response times were excluded from the relevant analyses. For math, test, and trait anxiety, we see an engagement bias for negative trials and a disengagement bias for math trials. For math and test anxiety, we observe disengagement bias for math trials. However, increases in trait anxiety are not associated with a significant change in disengagement bias for math trials.

Given the overlapping relationship between math anxiety and test anxiety, we evaluated how test anxiety influenced bias scores based on math and negative trials. We found a significant interaction, *F*(1,37) = 6.45, *p* = .015, *η*^2^ = .08. As test anxiety increased, individuals showed an increased engagement bias for negative trials. Conversely, as test anxiety increased, we see an increasing disengagement bias during math trials (Figure 2).

Although a similar relationship to test anxiety and math anxiety is observed in trait anxiety, this relationship is not statistically significant when trait (STAI) scores are used as a between-subject factor, *F*(1,36) = 3.05, *p* = .09, *η*^2^ = .01 (Figure 2). Indeed, as would be suggested by previous research on trait anxiety, higher scores on the STAI do seem to be associated with an engagement bias for negative stimuli, but increased trait anxiety results in no change in response to math trials.

In order to further examine how math anxiety influences responses to these stimuli while accounting for the contributions of other types of anxiety, we also calculated individual differences in math anxiety while controlling for various other self-report measures, such as test and trait anxiety. In a mixed ANOVA examining the effects of math anxiety controlling for trait anxiety (between-subject factor) on responses to math and negative stimuli (within-subject factor), math anxiety scores were not significantly related to responses to negative and math stimuli (*p* > .3). In a mixed ANOVA examining the effect of math anxiety controlling for test anxiety on negative and math attentional bias scores, there were no significant main effects or interactions (*p* > .3).

Across these measures of math and test anxiety, the pattern of responses in these behavioral data suggests that increased anxiety is associated with increased attentional engagement with negative information, suggesting that increases in anxiety are associated with increased orienting to negative information. Conversely, increased math and test anxiety (but not trait) are associated with attentional disengagement during math trials, suggesting that more math and test anxious individuals divert attention away from math compared to those low in anxiety.

### Imaging Results: Functional ROIs

We investigated whether amygdala reactivity elicited by brief stimulus presentation of math stimuli was correlated with math anxiety during presentations of mathematical stimuli. Using linear regression, we found that MARS score was not significantly related to amygdala reactivity, *p* = .18 uncorrected, *p* > .3, corrected. Comparing across LMA and HMA groups, we see increased amygdala reactivity in HMA compared to LMA participants, though the difference in activity was not statistically significant, *p* = .18 uncorrected, *p* > .3, corrected.

Signal change in the right amygdala was predicted by test anxiety, such that as TAI scores increase, activity in the right amygdala increases during math trials, *F*(1,37) = 4.15, *p* = .045 uncorrected, *p* < .10 corrected.

Amygdala responsivity to math during math trials was not significantly associated with trait anxiety, *p* > .3.

To further examine this relationship with math anxiety, we examined how math anxiety would influence amygdala reactivity, while controlling for trait and test anxiety. We found that when controlling for trait anxiety (MARS controlling for STAI), math anxiety is associated with increased amygdala activity during math trials, *F*(1,36) = 6.37, *p* = .02 uncorrected, *p* < .05 corrected, adjusted *R*^2^ = .13 (Figure 3). As math anxiety increases, activity in the right amygdala increases during trials where subjects are viewing mathematical expressions.

**Figure 3.**
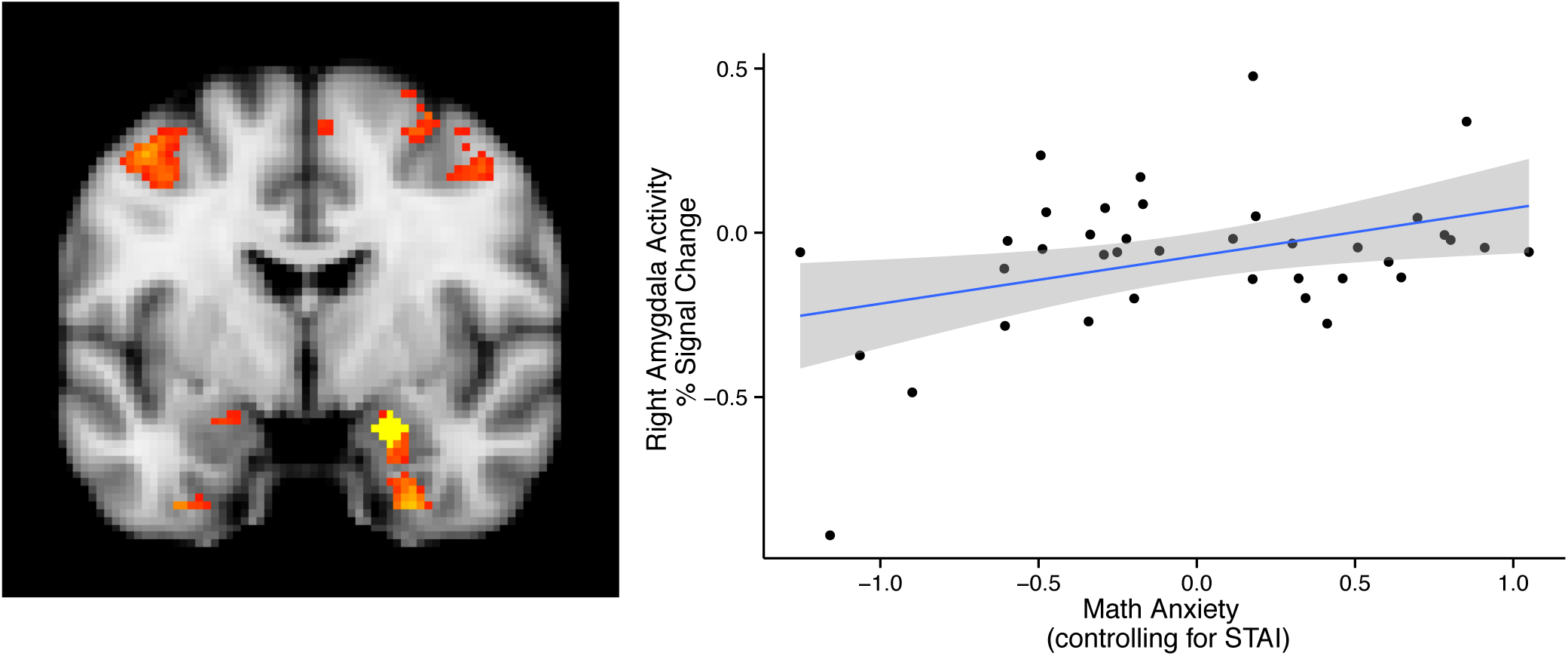
Right amygdala activity during math trials is positively correlated with math. Note: Percent signal change values (right) were extracted from the Math > Symbol contrast (Negative > Neutral depicted on the MNI brain template, left) in a spherical region of the amygdala isolated as a functional ROI centered on the peak voxel extracted from the group-level Negative > Neutral contrast (yellow ROI, left).

When controlling for test anxiety, math anxiety is not significantly correlated with amygdala reactivity during math trials, *p* = .99 uncorrected, *p* > .3 corrected.

Across measurements of math and test anxiety, we observe similar relationships that suggest increased anxiety is associated with increased amygdala reactivity when math is presented. These results are consistent with previous work on trait anxiety that associates increased anxiety with increased amygdala reactivity to negative cues (Bishop, 2004; Whalen, 2004a). Similar to what has been reported in specific phobia research (Sabatinelli, Bradley, Fitzsimmons, & Lang, 2005; Schienle et al., 2005), these results indicate increased amygdala activity related to a specific type of threat-related stimulus: math. This increase in amygdala response during math trials seems to be specific to math and test anxiety, as we do not observe the same relationship for trait anxiety.

### Anatomical ROI Analysis

Although we find the hypothesized relationship between math anxiety and amygdala activity in the functionally-defined (Negative >Neutral) ROI, it is unclear from the ROI analysis alone if this effect is specific to this sub-region of the amygdala. We find the same relationship identified by the functionally defined ROI: as math anxiety (controlling for trait anxiety) increases, amygdala activity is increased during presentations of mathematical stimuli, *F*(1,37) = 4.96, *p* = .03 uncorrected, adjusted *R*^2^= .09 (Figure 4). This relationship was not significantly associated with test anxiety, *p* = .18 uncorrected, math anxiety controlling for test anxiety, *p* = .99 uncorrected, or trait anxiety, *p* = .11 uncorrected. Thus, across methods of functional and anatomical ROI localization, we find the same relationship between math anxiety and increased amygdala activity when individuals are briefly exposed to mathematical stimuli in the dot probe task.

**Figure 4.**
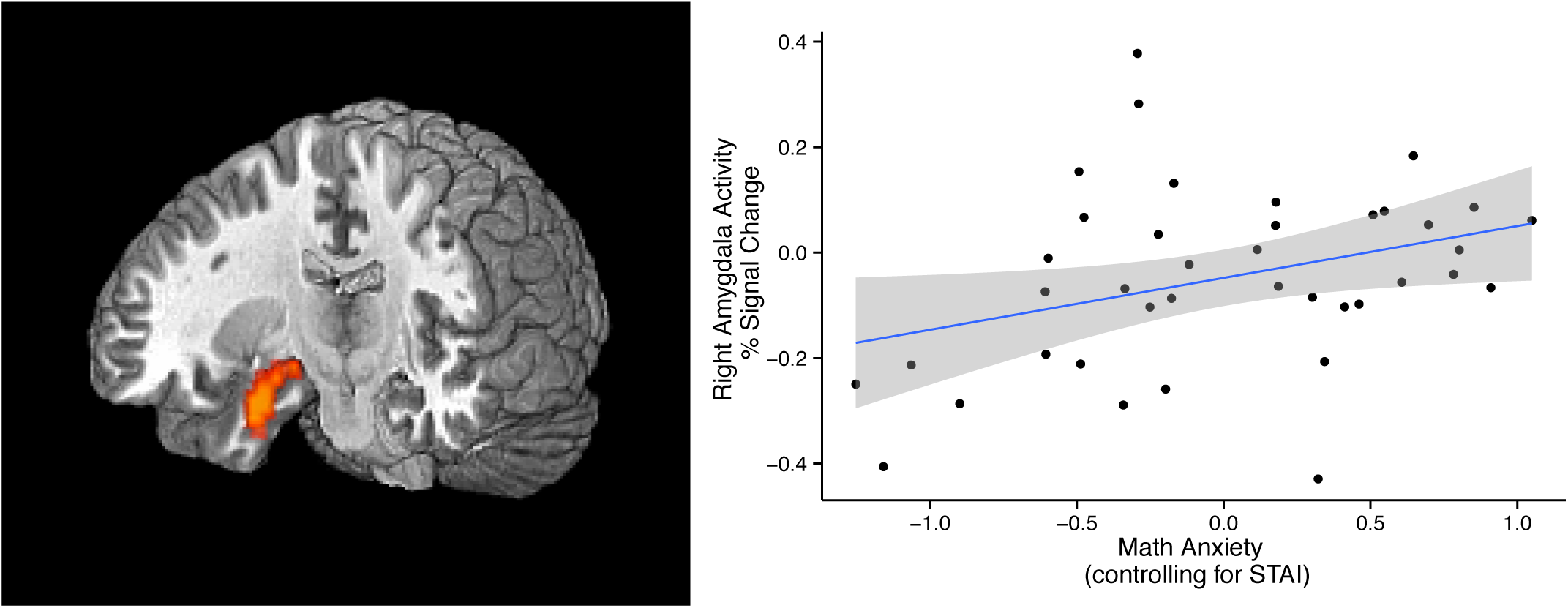
Anatomically-defined right amygdala activity during math trials is positively associated with math anxiety. Note. Math anxiety (controlling for trait anxiety) shows a similar correlation with the BOLD signal in the anatomically-defined right amygdala ROI (red, left).

### Whole Brain Analysis

To confirm the correlation between math anxiety and activity in the right amygdala during math trials observed in the functional ROI analysis, we conducted a whole-brain GLM analysis including MARS scores (controlling for STAI scores) as a parametric regressor. Among other regions activated by the presentation of math stimuli (Math > Symbol; Supplementary Table 2), we again find a cluster of activity in the right amygdala (*t*_*peak*_ = 2.77, MNI coordinates: *x* = 22, *y* = 8, *z* = -24). Greater math anxiety (controlling for trait anxiety) was associated with greater activity in the right amygdala, and yielded activations across the brain associated with motor execution, vision, and other cognitive processes. This consistency across methods of analysis provides further evidence for the idea that math stimuli are processed as though they are negatively-valenced stimuli for HMA individuals.

## Discussion

HMA individuals show increased amygdala reactivity in response to brief exposure to mathematical stimuli, illustrating that even without the anticipation or effortful computation associated with solving difficult mathematical problems, HMA individuals show a neural response related to threat and vigilance. This result is novel, in that it demonstrates that for HMA individuals, viewing mathematics may be similar to viewing a fear-conditioned stimulus, resulting in increased amygdala reactivity following mere exposure to the stimulus. In addition, HMA individuals show a behavioral disengagement bias specifically away from mathematical stimuli, compared to an engagement bias toward negatively-valenced images. This pattern of avoidance in math anxiety is similar to avoidance present in specific phobia (Pflugshaupt et al., 2005). Negative information still results in initial vigilance, as we see with the engagement bias for negative pictures (cf. Mogg, Bradley, & Williams, 1995). However, phobia-relevant stimuli may encourage subsequent avoidant reactions after initial orienting, as seen in the disengagement bias and amygdala reactivity for math stimuli. For HMA individuals, math stimuli appear to take on threat-like qualities.

As a way to better understand the characteristics of math anxiety, comparisons to other types of anxiety and to other patterns of neural reactivity and avoidant behaviors are informative, such as general or trait anxiety, and phobia. On the neural level, investigations utilizing fMRI have implicated the amygdala in attentional processing of threat-related information in anxious individuals (Bishop, 2008). For example, anxious individuals show increased amygdala reactivity to fear faces compared to non-anxious individuals even when the faces are not consciously perceived (Etkin et al., 2004), and when the fearful faces are irrelevant to the instructed task (Bishop, 2004). This finding bolsters the idea that attentional control mechanisms are altered in cases of increased anxiety, showing preferential reactivity to negative stimuli. Similarly, phobic individuals show increased amygdala reactivity specifically toward disorder-relevant stimuli compared to non-phobic individuals. For both snake (Sabatinelli et al., 2005) and spider phobia (Schienle et al., 2005), negative stimuli are associated with increased amygdala reactivity that is specific to the particular type of phobia. Similarly, our results indicate that increased math anxiety is associated with increased amygdala reactivity, specific to viewing mathematics (compared to other symbols).

While results indicating increased amygdala reactivity for mathematical stimuli provide strong support for the negative emotion at the root of math anxiety, based on previous research it could be consistent either with a characterization of math anxiety that resembles trait anxiety or specific phobia. Viewing these neural results in light of the behavioral pattern of attentional engagement or disengagement further distinguishes between these characterizations.

Previous research has characterized math anxiety in ways that are similar to trait anxiety, including the notion that math-related deficits result from changes in working memory due to increased rumination and negative self-talk (Beilock, 2008; Beilock & Ramirez, 2011). Trait anxiety has also been associated with disruptions in cognitive control and attentional orienting, generally characterized by a lower attentional threshold for negative information (Bishop, 2008; MacLeod & Mathews, 1988). Previous research using negatively-valenced stimuli has illustrated that increased anxiety is associated with an engagement bias for negative information (Bradley, et al., 1998; Mogg et al., 1995; Yiend & Mathews, 2010). We were able to replicate these findings when participants were shown negative stimuli, such that increased anxiety was associated with an engagement bias for negative stimuli (Figure 2). This is especially evident for trait anxiety, as increased scores on the STAI were associated with increased attentional bias for negative information, but no difference in attentional bias for math trials.

However, although we were able to replicate previous research using negative stimuli, the behavioral and neural results of this study indicate that math anxiety is instead more similar to state anxiety or to specific phobias in its behavioral phenotype characterized by avoidance of math. Previous studies have suggested that individuals with specific phobia (e.g., phobia of snakes) demonstrate an initial attentional orienting and increased amygdala reactivity toward phobia-related stimuli (Sabatinelli et al., 2005), but ultimately show a pattern of behavioral avoidance (Pflugshaupt et al., 2005). Applied to the context of math anxiety, in which math equations are the phobic stimuli, we observed that HMA participants to have an exaggerated reaction to math stimuli compared LMA individuals, as observed in increased amygdala reactivity, ultimately resulting in avoidance, or attentional disengagement.

A recent publication utilized a behavioral version of the dot probe task to assess whether this engagement bias seen for negative stimuli also exists for mathematical stimuli (Rubinsten, Eidlin, Wohl, & Akibli, 2015). In this attentional task, high math anxious individuals were found to have an engagement bias for math-related stimuli, and low math anxious individuals did not demonstrate this preferential processing. Our results did not replicate a previous investigation into math anxiety that found HMA individuals showed an engagement bias for math-related words and mathematical computations. However, the task in this prior study involved working memory-intensive strategies and may not have captured the initial, automatic orienting behaviors that might be a significant component of math anxiety. It is plausible that HMA individuals demonstrated an engagement bias due to the task difficulty, or due to differences in working memory capacity rather than differences in attentional deployment.

In this way, our paradigm, which allows participants to deploy attention to alternative stimuli, is a more valid depiction of how HMA individuals allocate attentional resources even when merely exposed to numerical or mathematical expressions. Further, that our work utilized fMRI to illustrate increased vigilance and threat-related amygdala processing even without requiring mathematical computations represents a significant improvement in our understanding of how anxious attention influences initial processing of math stimuli in HMA individuals. Perhaps future work will further investigate the interplay between attentional allocation and deployment of working memory resources, illustrating how both of these mechanism contribute to altered numerical computations in math anxiety.

The results of the present fMRI experiment illustrate that math anxiety is associated with changes in initial affective neural processing for math information, such that math stimuli – stimuli that would normally be considered to be affectively neutral stimuli – are rapidly processed like negative cues. Our results are consistent with previous research that also found that high math anxious children showed increased activity in the right amygdala while performing mathematical computations (Young et al., 2012). The present work extends this finding beyond the scope of children doing arithmetic math computations, as these results illustrate that even the mere presentation of mathematical expressions, without ever asking these participants to solve the expressions, is sufficient to cause increased amygdala reactivity in HMA young adults. In other words, for HMA individuals, math symbols might take on threat-like properties, and thus become associated with increased amygdala reactivity. These neural data suggest that HMA individuals have differential attentional mechanisms for math-related information, as in the current study even mere exposure to mathematical expressions was associated with increased amygdala reactivity for HMA individuals, as well as an attentional disengagement bias, diverting attention away from mathematical information. This exaggerated response to math was not observed in LMA individuals.

### Summary

The results of this study illustrate that for HMA individuals, exposure to mathematical stimuli results in aversive, distancing behavior and increased threat-related amygdala reactivity as anxious individuals disengage their attention from these threatening stimuli, similar to the response of phobic individuals to a phobic stimulus. This vigilance and disengagement occurs even when these individuals do not anticipate having to solve the problems, illustrating that these aversive reactions occur rapidly, and automatically. That these attentional biases influence low-level cognitive and neural processes has significance for how we conceptualize treatments for math anxiety. This disengagement bias speaks to a broader pattern of avoidance and to the aversive nature of math for HMA individuals. In the long-term, math anxious individuals show avoidance of math computations (e.g., speed-accuracy tradeoffs; (Faust, 1996), choose not to take advanced math classes, and choose careers that do not involve math (Ashcraft, 2002). The present results add to this picture a rapid, short-term attentional disengagement from mathematical information that is also associated with math anxiety.

Further investigations into the neural changes associated with math anxiety might investigate how individual differences in math anxiety influence the recruitment of regions associated with cognitive and affective control, as these neural results indicate that individuals may need to overcome initial neural signals of increased vigilance in order to engage regions of the brain associated with mathematical computations. Future interventions might target improved attentional engagement and approach-related behavior (similar to systematic desensitization, a common treatment for phobia) when individuals are exposed to math and asked to solve math problems. By improving how we conceptualize the cognitive and neural bases of math anxiety, we can continue to understand how specific types of anxiety – or phobias – influence cognition, and thus achieve further insight into improving performance deficits caused by math anxiety.

## References

Ashcraft, M. H. (2002). Math anxiety: Personal, educational, and cognitive consequences. Current Directions in Psychological Science, 11(5), 181–185.

Beilock, S. L. (2008). Math performance in stressful situations. Current Directions in Psychological Science, 17(5), 339–343.

Beilock, S. L., & Ramirez, G. (2011). 5 On the Interplay of Emotion and Cognitive Control: Implications for Enhancing Academic Achievement. Psychology of Learning and Motivation, 55, 137. http://doi.org/10.1016/B978-0-12-387691-1.X0001-4

Bishop, S. J. (2004). State Anxiety Modulation of the Amygdala Response to Unattended Threat-Related Stimuli. Journal of Neuroscience, 24(46), 10364–10368. http://doi.org/10.1523/JNEUROSCI.2550-04.2004

Bishop, S. J. (2007). Neurocognitive mechanisms of anxiety: an integrative account. Trends in Cognitive Sciences, 11(7), 307–316. http://doi.org/10.1016/j.tics.2007.05.008

Bishop, S. J. (2008). Neural Mechanisms Underlying Selective Attention to Threat. Annals of the New York Academy of Sciences, 1129(1), 141–152. http://doi.org/10.1196/annals.1417.016

Bradley, B. P., Mogg, K., Falla, S. J., & Hamilton, L. R. (1998). Attentional Bias for Threatening Facial Expressions in Anxiety: Manipulation of Stimulus Duration. Cognition & Emotion, 12(6), 737–753. http://doi.org/10.1080/026999398379411

Dale, A. M., Fischl, B., & Sereno, M. I. (1999). Cortical Surface-Based Analysis. NeuroImage, 9(2), 179–194. http://doi.org/10.1006/nimg.1998.0395

Daly, J. A., & Wilson, D. A. (1983). Writing apprehension, self-esteem, and personality. Research in the Teaching of English, 327–341.

Davis, M., & Whalen, P. J. (2001). The amygdala: vigilance and emotion. Molecular Psychiatry,6(1), 13–34.

Dehaene, S., Spelke, E., Pinel, P., Stanescu, R., & Tsivkin, S. (1999). Sources of mathematical thinking: behavioral and brain-imaging evidence. Science (New York, N.Y.), 284(5416), 970– 974. http://doi.org/10.2307/25464011?ref=search-gateway:e3793a486fd474eb3d745fd7e7867320

Etkin, A., Klemenhagen, K. C., Dudman, J. T., Rogan, M. T., Hen, R., Kandel, E. R., & Hirsch, J. (2004). Individual differences in trait anxiety predict the response of the basolateral amygdala to unconsciously processed fearful faces. Neuron, 44(6), 1043–1055. http://doi.org/10.1016/j.neuron.2004.12.006

Faust, M. W. (1996). Mathematics Anxiety Effects in Simple and Complex Addition. Mathematical Cognition, 2(1), 25–62. http://doi.org/10.1080/135467996387534

Hembree, R. (1990). The nature, effects, and relief of mathematics anxiety. Journal for Research in Mathematics Education, 33–46.

Holland, P. C., & Gallagher, M. (1999). Amygdala circuitry in attentional and representational processes. Trends in Cognitive Sciences, 3(2), 65–73.

Lang, P., & Bradley, M. M. (2007). The International Affective Picture System (IAPS) in the study of emotion and attention. Handbook of Emotion Elicitation and ….

Lyons, I. M., & Beilock, S. L. (2012a). Mathematics Anxiety: Separating the Math from the Anxiety. Cerebral Cortex, 22(9), 2102–2110. http://doi.org/10.1093/cercor/bhr289

Lyons, I. M., & Beilock, S. L. (2012b). When Math Hurts: Math Anxiety Predicts Pain Network Activation in Anticipation of Doing Math. PLoS ONE, 7(10), e48076. http://doi.org/10.1371/journal.pone.0048076.t003

MacLeod, C., & Mathews, A. (1988). Anxiety and the allocation of attention to threat. The Quarterly Journal of Experimental Psychology Section A, 40(4), 653–670. http://doi.org/10.1080/14640748808402292

Mogg, K., Bradley, B. P., & Williams, R. (1995). Attentional bias in anxiety and depression: The role of awareness. British Journal of Clinical Psychology, 34(1), 17–36. http://doi.org/10.1111/j.2044-8260.1995.tb01434.x

Pflugshaupt, T., Mosimann, U. P., Wartburg, R. V., Schmitt, W., Nyffeler, T., & Müri, R. M. (2005). Hypervigilance–avoidance pattern in spider phobia. Journal of Anxiety Disorders, 19(1), 105–116. http://doi.org/10.1016/j.janxdis.2003.12.002

Rubinsten, O., Eidlin, H., Wohl, H., & Akibli, O. (2015). Attentional bias in math anxiety. Frontiers in Psychology, 6, 1539–1539. http://doi.org/10.3389/fpsyg.2015.01539

Sabatinelli, D., Bradley, M. M., Fitzsimmons, J. R., & Lang, P. J. (2005). Parallel amygdala and inferotemporal activation reflect emotional intensity and fear relevance. NeuroImage, 24(4), 1265–1270. http://doi.org/10.1016/j.neuroimage.2004.12.015

Schienle, A., Schäfer, A., Walter, B., Stark, R., & Vaitl, D. (2005). Brain activation of spider phobics towards disorder-relevant, generally disgust- and fear-inducing pictures. Neuroscience Letters, 388(1), 1–6. http://doi.org/10.1016/j.neulet.2005.06.025

Spielberger, C. D. (2009). Test anxiety inventory. In I. B. Weiner & W. E. Craighead (Eds.), The Corsini Encyclopedia of Psychology. John Wiley & Sons, Inc.

Spielberger, C. D. (2010). State-Trait Anxiety Inventory. In The Corsini Encyclopedia of Psychology. Hoboken, NJ, USA: John Wiley & Sons, Inc. http://doi.org/10.1002/9780470479216.corpsy0943

Suinn, R. M., & Winston, E. H. (2003). The Mathematics Anxiety Rating Scale, a brief version: psychometric data. Psychological Reports, 92(1), 167–173.

Watson, D., Clark, L. A., & Tellegen, A. (1988). Development and validation of brief measures of positive and negative affect: the PANAS scales. Journal of Personality and Social Psychology, 54(6), 1063–1070. http://doi.org/10.1037/0022-3514.54.6.1063

Whalen, P. J. (1998). Fear, vigilance, and ambiguity: Initial neuroimaging studies of the human amygdala. Current Directions in Psychological Science, 7(6), 177–188.

Whalen, P. J. (2004a). Fear, vigilance, and ambiguity: Initial neuroimaging studies of the human amygdala, 1–12.

Whalen, P. J. (2004b). Human Amygdala Responsivity to Masked Fearful Eye Whites. Science (New York, N.Y.), 306(5704), 2061–2061. http://doi.org/10.1126/science.1103617

Yiend, J., & Mathews, A. (2010). Anxiety and attention to threatening pictures. The Quarterly Journal of Experimental Psychology Section A, 54(3), 665–681. http://doi.org/10.1080/713755991

Young, C. B., Wu, S. S., & Menon, V. (2012). The neurodevelopmental basis of math anxiety. Psychological Science, 23(5), 492–501. http://doi.org/10.1177/0956797611429134

